# Shared characteristics underpinning C_4_ leaf maturation derived from analysis of multiple C_3_ and C_4_ species of *Flaveria*

**DOI:** 10.1101/088153

**Authors:** Britta M.C. Kümpers, Steven J. Burgess, Ivan Reyna-Llorens, Richard Smith-Unna, Chris Boursnell, Julian M. Hibberd

**Affiliations:** Department of Plant Sciences, Downing Street, University of Cambridge, Cambridge CB2 3EA, UK

**Keywords:** C_4_ photosynthesis, C_4_ leaf anatomy, parallel evolution, convergent evolution, gene expression, RNA-SEQ, *Flaveria*

## Abstract

Most terrestrial plants use C_3_ photosynthesis to fix carbon. In multiple plant lineages a modified system known as C_4_ photosynthesis has evolved. To better understand the molecular patterns associated with induction of C_4_ photosynthesis the genus *Flaveria* that contains C_3_ and C_4_ species was used. A base to tip maturation gradient of leaf anatomy was defined, and RNA sequencing was undertaken along this gradient for two C_3_ and two C_4_ *Flaveria* species. Key C_4_ traits including vein density, mesophyll and bundle sheath cross-sectional area, chloroplast ultrastructure, and abundance of transcripts encoding proteins of C_4_ photosynthesis were quantified. Candidate genes underlying each of these C_4_ characteristics were identified. Principal Components Analysis indicated that leaf maturation and then photosynthetic pathway were responsible for the greatest amount of variation in transcript abundance. Photosynthesis genes were over-represented for a prolonged period in the C_4_ species. Through comparison with publically available datasets we identify a small number of transcriptional regulators that have been up-regulated in diverse C_4_ species. The analysis identifies similar patterns of expression in independent C_4_ lineages and so indicates that the complex C_4_ pathway is associated with parallel as well as convergent evolution.

**Highlight:** We identify transcription factors that show conserved patterns of expression in multiple 29 C_4_ species, both within the Flaveria genus, but also in more distantly related C_4_ plants.

## Introduction

Photosynthesis powers life on earth by harvesting energy from sunlight to fix carbon dioxide. In seed plants three types of photosynthesis known as C_3_, C_4_ and Crassulacean Acid Metabolism (CAM) have been defined. C_3_ photosynthesis is considered the ancestral form and is found in the majority of species. All three photosynthetic pathways use Ribulose-1,5-Bisphosphate Carboxylase Oxygenase (RuBisCO) to catalyse CO_2_ fixation in the Calvin-Benson-Bassham (CBB) cycle, and in so doing, two molecules of the three-carbon molecule 3-phosphoglycerate are formed. However, RuBisCO is not completely substrate specific, catalysing an oxygenation reaction that produces 2-phosphoglycolate as well as 3-phosphoglycerate (Bowes *et al.*, 1971). As phosphoglycolate is toxic and removes carbon from the CBB cycle, it is rapidly metabolised by the photorespiratory pathway to ensure it does not accumulate, but also to retrieve carbon. Photorespiration uses energy inputs and is not completely effective in terms of carbon retrieval so some CO_2_ is lost (Bauwe *et al.*, 2010). The rate of oxygenation at the active site of RuBisCO increases with temperature, and so in lower latitudes, multiple plants linages have evolved either CAM or C_4_ photosynthesis, which in both cases, initially fix CO_2_ by an alternate carboxylase, and then subsequently release high concentrations of CO_2_ around RuBisCO to limit oxygenation.

Almost all C_4_ plants use a two-celled system in which Phospho*enol*pyruvate Carboxylase (PEPC) initially fixes carbon in Mesophyll (M) cells, and then after release of CO_2_ in the Bundle Sheath (BS) cells, RuBisCO re-fixes CO_2_ in the CBB cycle. The production of CO_2_ within the BS leads to high concentrations of CO_2_ around RuBisCO and minimises oxygenation. PEPC generates oxaloacetate, which is rapidly metabolised to malate and/or aspartate prior to their diffusion to the BS. Three C_4_ acid decarboxylases enzymes known as NADP-malic enzyme (NADP-ME), NAD-malic enzyme (NAD-ME) and Phospho*enol*pyruvate Carboxykinase (PEPCK) have been shown to release CO_2_ around RuBisCO, and for many years, based on their relative abundance, these decarboxylase enzymes were used to define three C_4_ biochemical subtypes. More recently it has been shown that this division into three types is less rigid than previously (Furbank, 2011; Bellasio and Griffiths, 2014; Wang *et al.*, 2014). The variation in the activity of these three decarboxylases indicates that C_4_ photosynthesis is underpinned by convergent evolution in different lineages of plants. Within the more than sixty groups of plants known to use the C_4_ pathway, there are some genera containing both C_3_ and C_4_ species (Sage *et al.*, 2011). Probably, the most studied of these clades with closely related C_3_ and C_4_ species is the genus *Flaveria* found in the Asteraceae (Drincovich *et al.*, 1998; Gowik *et al.*, 2011; Schulze *et al.*, 2013).

Most species of *Flaveria* are native to Central and North America and grow as annual or perennial herbs or shrubs with decussate leaves (Powell, 1978). C_4_ species of *Flaveria* show high activities of the NADP-ME C_4_ acid decarboxylase in chloroplasts of the BS, and leaf anatomy conforms to the Atriplicoid type (McKown and Dengler, 2007). Phylogenetic reconstruction of this group has been conducted using morphological as well as life history and gene sequence data for 21 of the 23 known species (McKown *et al.*, 2005; Lyu *et al.*, 2015). The consensus from this work is that the ancestral condition in *Flaveria* is C_3_ photosynthesis (McKown *et al.*, 2005; Lyu *et al.*, 2015) and that the occurrence of multiple C_3_ and C_4_ species within the same genus provides an interesting system to study processes associated with the C_4_ phenotype.

In this study, we used two pairs of C_3_ and C_4_ *Flaveria* species, and set out to link the gradual maturation of C_3_ and C_4_ characteristics in leaves to underlying alterations in transcript abundance. By linking the development of the C_4_ phenotype to changes in gene expression in two C_4_ species, and comparing these findings with equivalent data from two C_3_ species in the same genus, we aimed to identify common traits associated with C_4_ photosynthesis, and to remove species-specific characteristics from our datasets. Using the maturing leaf as a dynamic system we show that in both C_4_ species studied, the induction of Kranz anatomy occurs along a base to tip developmental gradient in leaves under 2cm length. We sampled this maturation gradient and undertook RNA sequencing to correlate the underlying patterns of gene expression with anatomical development.

## Materials & Methods

### Plant growth

*Flaveria bidentis* (L.) Kuntze, *Flaveria pringlei* Gandoger, *Flaveria robusta* Rose, and *Flaveria trinervia* (Spreng.) C. Mohr were grown in a glasshouse on the roof of the Department of Plant Sciences, Cambridge. Temperature was maintained above 20°C and supplemental lighting provided to ensure at least 250-350 μmol m^−2^ s^−1^ photon flux density for 16 hours per day. Seeds were sown directly onto soil (Levington’s M3 potting compost; Scotts Miracle-Gro Company, Godalming, UK) in covered pots as seeds need high humidity for germination. Covers were removed when seedlings were 1 cm high.

### Analysis of leaf anatomy

Samples of 4 mm^2^ to 1 cm^2^ were fixed in 4% (w/v) formaldehyde at 4°C overnight and then placed on ice and dehydrated prior to being placed in 100% (v/v) ethanol, followed by 1:1 ethanol/Technovit mix and then 100% Technovit 7100 (Heraeus Kulzer, Germany). Samples were subsequently left in Technovit solution plus hardener I (1g hardener per 100ml) for at least an hour. Disposable plastic resin embedding moulds (Agar Scientific, UK) were filled with a mixture of Technovit plus hardener I and II (15 ml of Technovit plus hardener I were mixed with 1 ml of hardener II). Samples were arranged within the embedding moulds which were then covered with unstretched Parafilm^®^ M to seal them from air, and left to harden overnight. Samples were removed, heated to 80°C and trimmed for sectioning. Sections of 2μm thickness were produced using a Thermo Scientific Microm HM340E microtome. Ribbons were mounted onto SuperFrost®white microscope slides (VWR, Leuven, NL), left to dry and then stained with 0.1% (w/v) toluidine blue. All sections were analysed with a BX41 light microscope (Olympus, Center Valley, PA, USA), usually using the bright-field setting.

To clear leaves, they were placed into 70% (v/v) ethanol and heated to 80°C. The next day samples were placed in 5% (w/v) NaOH for about 15 minutes to clear leaves further and then mounted in water and analysed by light microscopy. To quantify leaf anatomical characteristics, Photoshop CS5 was used. The programme was calibrated with scale bars and the lasso tool was used to measure cell area for both BS and M cells. Measurements of M cells were taken between BS cells, with 30 BS and 30 M cells being measured per leaf section. This was done for all six leaf stages and for three biological replicates for each species. Images derived from leaf sections were assembled using Photoshop. The background of sections was averaged and in some cases the contrast was increased to improve visibility of cell borders. Vein density was determined using images of cleared leaves. In mature sections of the leaf there was no ambiguity in collecting these data, whereas in the most basal sections of some leaves, the estimates may be underestimates of the extent of venation as immature veins are not always visible. Three independent leaves were measured per species. The length of the veins was measured using Q-Capture Pro7 and vein density was expressed in mm mm^−2^.

Samples for the Transmission Electron Microscope (TEM) were processed by the Multi-Imaging Centre in the Department of Physiology, Development and Neuroscience (Cambridge). Sections of about 1 mm thickness of all six stages along a developing leaf were used for embedding. Tissues were fixed in in 4% (w/v) glutaraldehyde in 0.1 M HEPES buffer with a pH of 7.4 for 12 hours at 4°C. Subsequently they were rinsed in 0.1 M HEPES buffer five times, and then treated with 1% osmium ferricyanide at room temperature for two hours and rinsed in deionised water five times before being treated with 2% (w/v) uranyl acetate in 0.05 M maleate buffer with a pH of 5.5 for two hours at room temperature. They were rinsed again in deionised water and dehydrated in an ascending series of ethanol solutions from 70% to 100% (v/v). This was followed by treatment with two changes of dry acetonitrile and infiltration with Quetol 651 epoxy resin. 50-70 nm thick sections were cut with a Leica Ultracut UCT and stained with lead citrate and uranyl acetate. Images were taken with a Tecnai G2 TEM (FEI, Hillsboro, USA) and operated at 120Kv using an AMT XR60B digital camera (Advanced Microscopy Techniques, Woburn, USA) running Deben software (Deben UK Limited, Bury St. Edmunds, UK). Images were taken at 1700x, 3500x and 5000x magnification.

### RNA extraction and sequencing

Deep sequencing was carried out to analyse transcriptomes associated with the leaf developmental gradient defined by the anatomical analysis. Opposite leaves from the third leaf pair after the cotyledons were harvested in the morning between 10 am and 11am (four hours after the start of the photoperiod in the glasshouse). The leaves were harvested when they had reached 1.8-2 cm length (from base to tip). Leaves were measured on an ice-cold glass plate and cut into six portions of equal length (3-3.3 mm). Each sample was then immediately placed into liquid nitrogen. Sections of twelve leaves were collected before extraction of RNA using the Qiagen Plant RNeasy Mini Kit. The optional DNA digestion step with RNAse-free DNAse was performed. Three biological replicates were analysed for each of the four *Flaveria* species. Total leaf RNA was sent to the Genomics & Transcriptomics Laboratory (GTL) in Düsseldorf (Germany) for paired-end sequencing. RNA samples were prepared following the TrueSeq RNA sample preparation v2 guide (Revision F) and sequenced using an Illumina/Solexa HiSeq 2000 machine. *De-novo* assembly was undertaken following previously published methods (Aubry *et al.*, 2014; Smith-Unna *et al.*, 2016). For Principal Components Analysis Z-scores were calculated from Transcripts per Million (TPM) reads from for each gene across the four species. Expression patterns of genes of interest were analysed using a custom-made R script to extract the data of interest and to visualise expression patterns. GO term analysis was performed according to (Burgess *et al.*, 2016).

## Results and Discussion

### Leaf maturation in C_3_ and C_4_ *Flaveria* species

We first confirmed that fully expanded mature leaves of C_3_ *F. pringlei* and *F. robusta* as well as C_4_ *F. bidentis* and *F. trinervia* showed characteristic C_3_ and C_4_ anatomy under our growth conditions. In all cases the third leaf pair was chosen as the first and second leaf pairs have previously been described as juvenile (McKown and Dengler, 2009). Mature third leaves of C_3_ and C_4_ *Flaveria* ranged from 6-10 cm in length, but the C_3_ species had narrower leaves with an entire margin whereas the C_4_ leaves were wider and the margin was slightly dentate (Figure 1A). Both C_4_ species had closer veins than the C_3_ species (Figure 1B, Supplementary Figure 1). In addition, analysis of transverse leaf sections indicated that C_3_ leaves were thicker because of larger cells and more cell layers (Figure 1C). Clear differences in BS and M cell arrangement were visible between the C_3_ and C_4_ pairs with the BS in both C_4_ species being more uniform in shape than in the C_3_ species (Figure 1C). In addition, compared with the C_3_ species, M cells in the C_4_ leaves were smaller (Supplementary Figure 1) and showed increased contact with the BS. In agreement with previous analysis (McKown and Dengler, 2007), leaves of the two C_4_ and two C_3_ species typically contained five and eight ground cell layers between the adaxial and abaxial epidermis respectively.

**Figure 1:**
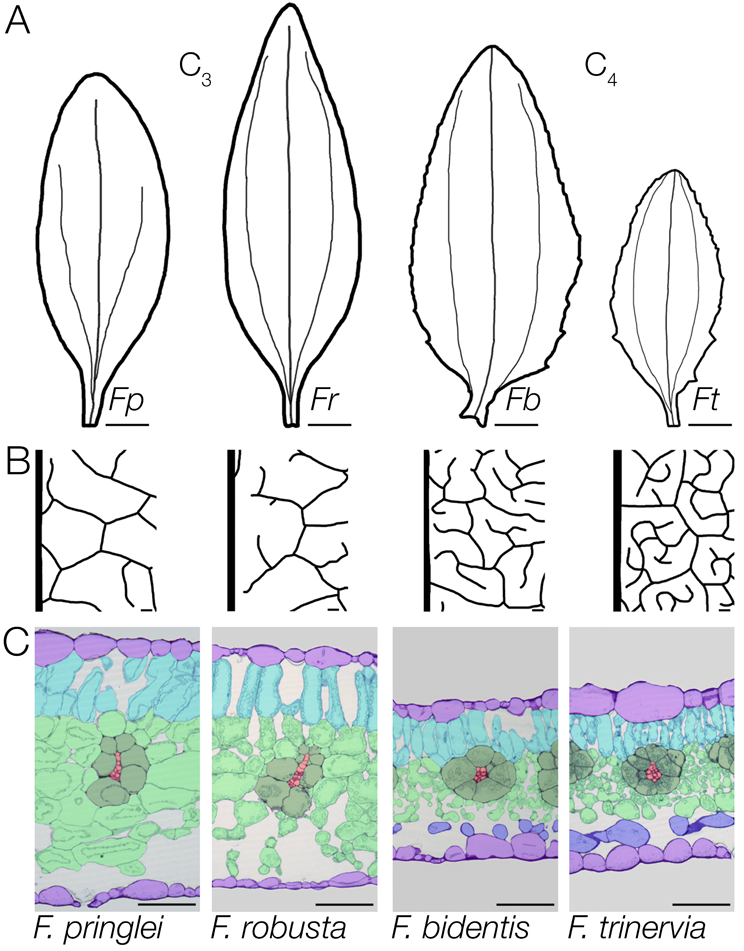
Characteristics of mature leaves from C_3_ and C_4_ species of *Flaveria*. **A**: Representative outlines of mature leaves showing the variation between species and the dentate nature of the two C_4_ species. **B**: Vein traces taken in the middle of the leaf to the right of the midvein indicating denser venation in mature leaves of both C_4_ species. **C**: Transverse sections showing the reduced mesophyll cross-sectional area and the closer veins with more symmetric bundle sheaths in the C_4_ compared with the C_3_ species. Each cell-type is false coloured-coded: veins (red), bundle sheath (dark green), spongy mesophyll (light green), palisade mesophyll (turquoise), parenchyma layer with no or few chloroplasts (blue) and epidermis (lilac). Species abbreviations are as follows: *Flaveria pringlei (Fp), Flaveria robusta (Fr), Flaveria bidentis (Fb), Flaveria trinervia, (Ft)*. Scale bars represent 1cm (**A**) or 100μm (**B-C**).

In all four *Flaveria* species, leaves of 1.8-2cm length showed a basipetal maturation gradient with differentiating tissues at the base and fully differentiated tissues at the tip (Figure 2A-C and Supplementary Figure 2). This base-to-tip maturation programme is typical of dicotyledons, but it is notable that in *Flaveria* maturation occurred in larger leaves than in *G. gynandra,* where the maturation gradient was detected in leaflets of 0.4 cm length (Aubry *et al.*, 2014). This prolonged development of leaf maturation in *Flaveria* thus provides an excellent system to analyse the induction of C_4_ photosynthesis as the larger leaf allows these leaves to be divided into more stages for functional analysis.

**Figure 2:**
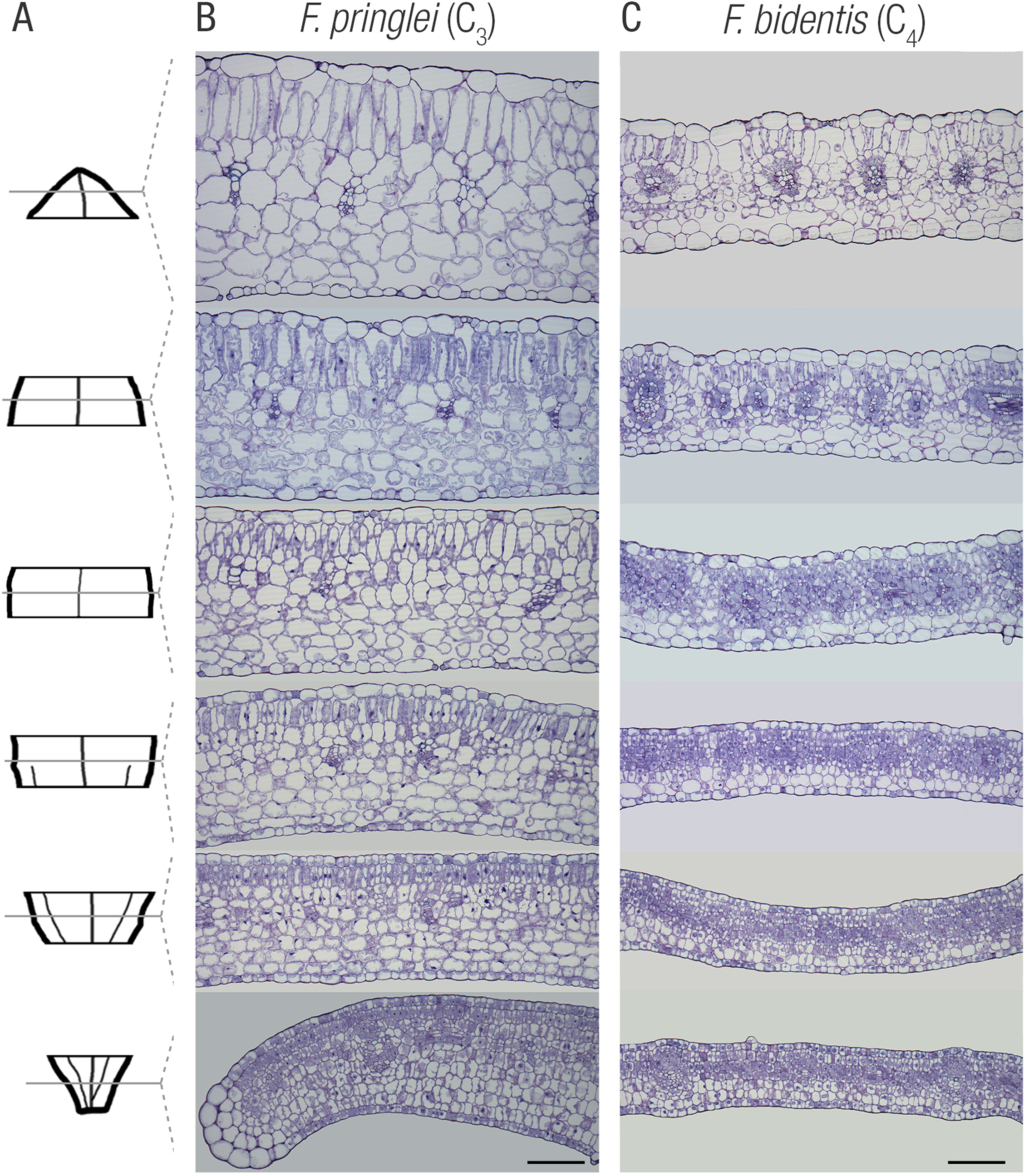
Within leaf sampling indicates the gradual maturation gradient in leaf anatomy for C_3_ and C_4_ *Flaveria* speciecs. **A**: Representative leaf outline illustrating leaf sampling. Representative transverse sections from C_3_ *Flaveria pringlei* (**B**) and C_4_ *Flaveria bidentis* (**C**) from base (bottom) to tip (top). Note the gradual expansion of cells, increased vacuolisation and clearer delineation of both mesophyll and bundle sheath cells from base to tip. Scale bars represent 100μm.

To better understand differences in leaf maturation in C_3_ and C_4_ *Flaveria*, leaves from each species were divided into six portions, RNA was extracted, quantified and integrity determined (Supplementary Figure 3) and then subjected to deep sequencing. On average 20 million reads were recovered from each replicate (Supplementary Table 1). After *de novo* assembly of these reads, the average number of annotated transcripts per species was 12,475 (Supplementary Table 1).

Read mapping was used to quantify transcript abundance in Transcripts Per Million (TPM, Supplementary Data 1). By grouping the two base, mid and tip stages, it was possible to quickly analyse transcript behaviour across the maturation gradient. This revealed that on average 40% of annotated transcripts showed descending behaviours, meaning that they were expressed more strongly at the base of the leaf than at the tip (Supplementary Table 1). Using the 8000 annotations found in all four species, correlation analysis showed that patterns of gene expression first clustered by species and then by photosynthetic type, meaning that the correlation between neighbouring developmental stages was highest within species and that the correlation was higher between species of the same photosynthetic type than between species of different photosynthetic types (Supplementary Figure 4). A gradient from base to tip was clear in all four species but was more pronounced in the two C_4_ species (Supplementary Figure 4).

The six portions from the base to the tip of the leaf showed a clear induction of genes known to encode components of the C_4_ cycle (Figure 3). Around 200 genes are consistently up-regulated in the upper tip tissue using C_4_ compared with C_3_ photosynthesis (Figure 4A, Supplementary Data 2), and in young tissue that is still undergoing differentiation, the number of genes up-regulated in the C_4_ species is around double this number. These estimates are significantly lower than those reported from analysis of whole leaves at various developmental stages from C_3_ *Tarenaya hassleriana* and C_4_ *Gynandropsis gynandra* in the Cleomaceae (Külahoglu *et al.*, 2014), where between 2500 and 3500 genes were estimated to be upregulated in the C_4_ species. This reduction in differential gene expression in the C_4_ compared with C_3_ *Flaverias* in this study compared with the estimates from the Cleomaceae may be due to the use of two C_3_ and two C_4_ species removing species-specific differences, but also to the reduced phylogenetic distance between the *Flaveria* species. Principal Components Analysis indicated that the first two dimensions contributed to 37 and 19% of the variation respectively, and that whereas the first dimension was associated with the leaf maturation signal from all species, the second dimension was associated with the photosynthetic pathway being used (Figure 4B). This finding implies that C_4_ metabolism has a large impact on the differences in gene expression associated with C_3_ and C_4_ leaves within one genus. When MapMan categories associated with the top decile of differentially expressed genes were assessed it was notable that photosynthesis genes were over-represented for a prolonged period along the gradient in the C_4_ species (Figure 4C). This was also the case for genes related to DNA and transport. In contrast, the MapMan category associated with RNA was over-represented for longer in the C_3_ species (Figure 4C). Equivalent analysis for Gene Ontology (GO) terms indicated that cell wall loosening was over-represented for longer in both C_4_ compared with the C_3_ species (Supplementary Figure 5). Overall, these data indicate that the maturation of *Flaveria* leaves captures changes in transcript abundance associated with induction of C_3_ or C_4_ photosynthesis, and provide insight into the broader alterations in gene expression. We next aimed to relate the underlying anatomical changes associated with leaf maturation in these C_3_ and C_4_ species of *Flaveria* to changes in gene expression determined from the deep sequencing.

**Figure 3:**
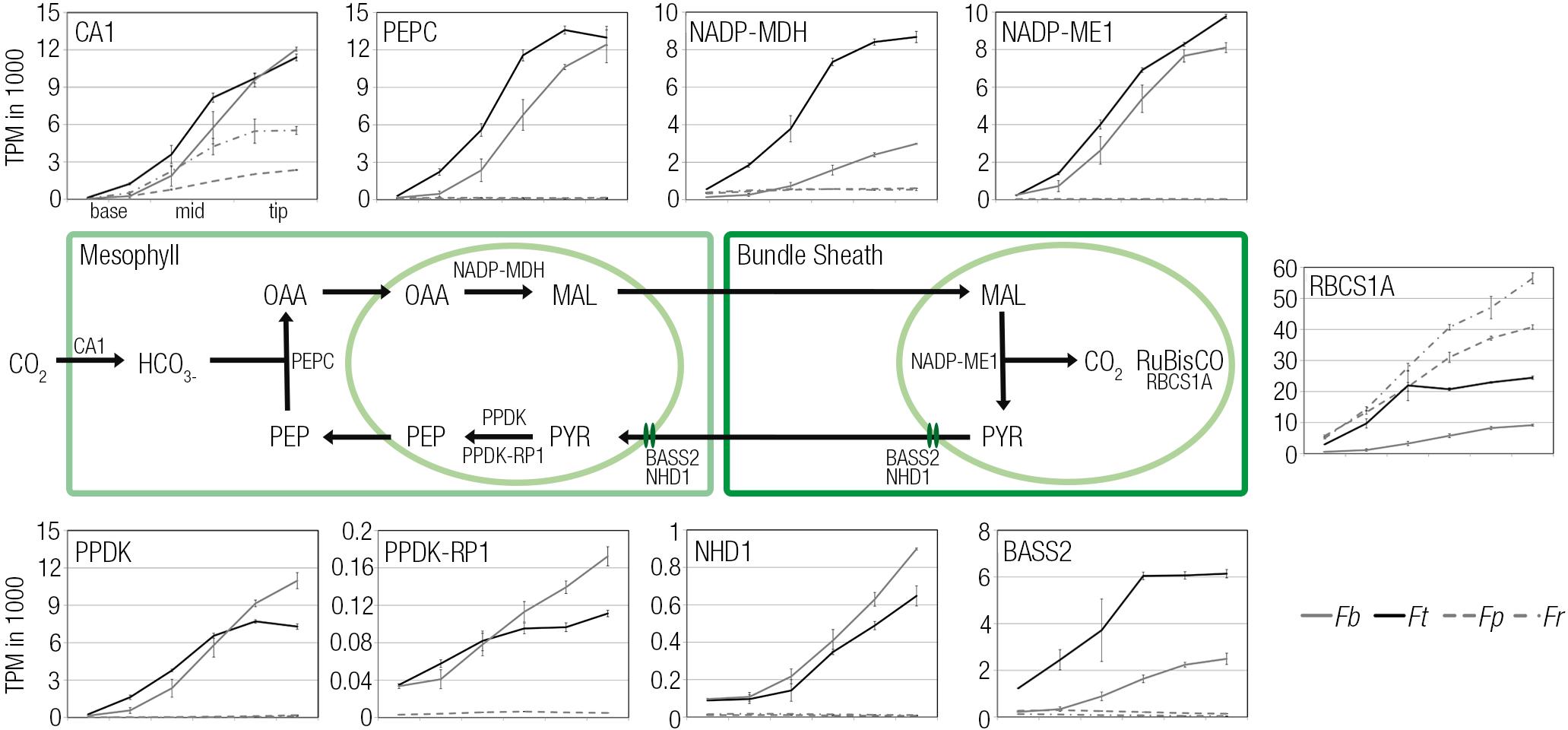
Transcripts encoding the C_4_ cycle increase dramatically during leaf maturation of the C_4_ species. The schematic of major components of the C_4_ pathway located in mesophyll and bundle sheath cells of C_4_ leaves is surrounded by plots of transcript abundance for key genes of the NADP-ME sub-type in transcripts per million (TPM) for the four species. *RBCS1A* transcripts are, as expected higher in the C_3_ species. All C_4_ transcripts other than *CARBONIC ANHYDRASE1* (*CA1*) are barely detectable in the C_3_ species. *PEPC1 = PHOSPHOENOLPYRUVATE CARBOXYLASE 1, NADP-MDH = NADP-DEPENDENT MALATE DEHYDROGENASE, NADP-ME1 = NADP-DEPENDENT MALIC ENZYME1, PPDK= PYRUVATE,ORTHOPHOSPHATE DIKINASE, PPDK-RP1 = PYRUVATE,ORRTHOPHOSPHATE DIKINASE REGULATORY PROTEIN, NHD1 = SODIUM:HYDROGEN ANTIPORTER, BASS2 = BILE ACID SODIUM SYMPORTER FAMILY2.* The two C_4_ species are shown with solid lines, and the two C_3_ species with dashed lines.

**Figure 4:**
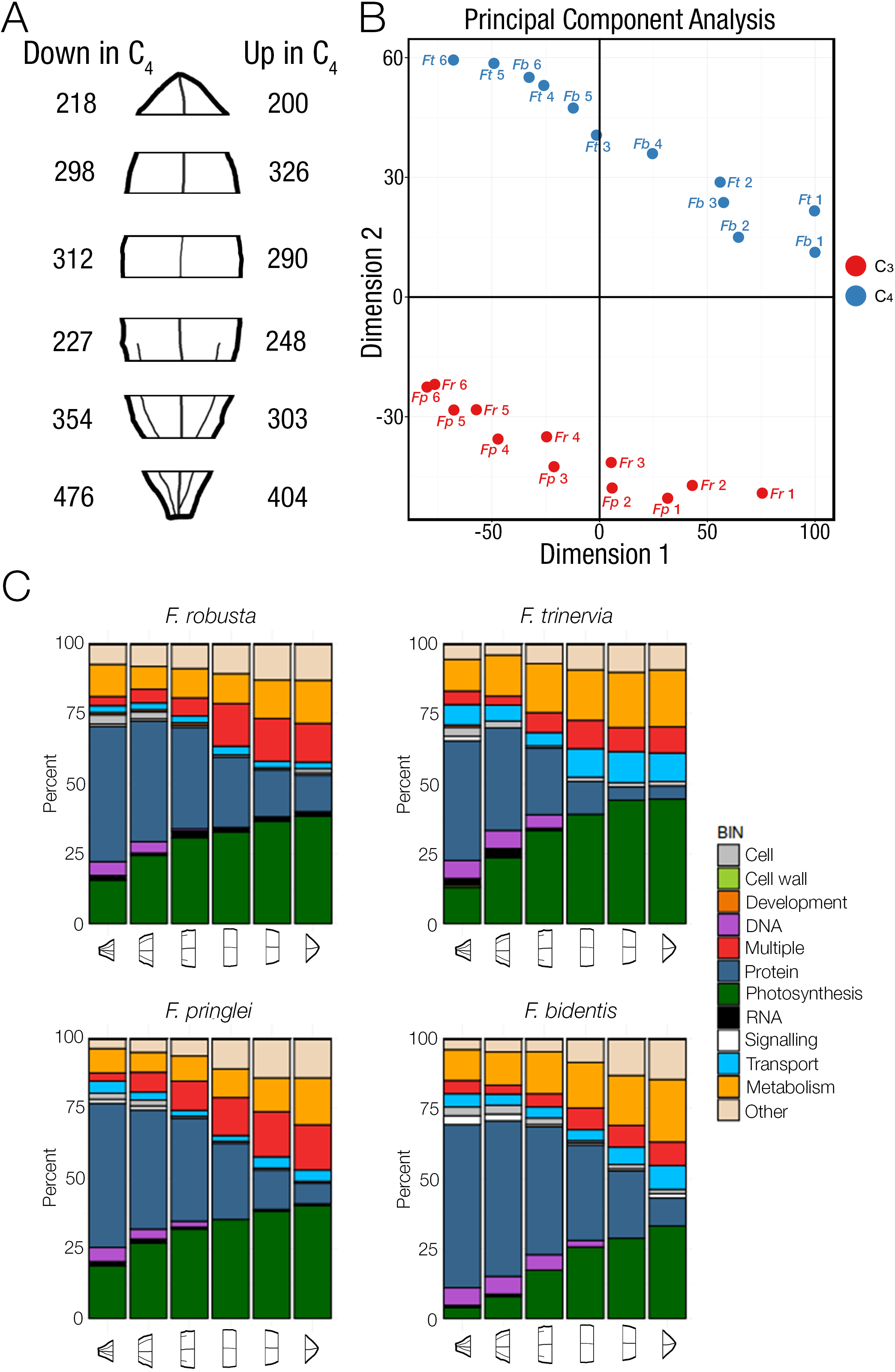
Overview of the global changes in gene expression along the C_3_ and C_4_ *Flaveria* maturation gradients. **A**. RNA-seq was conducted on six regions of *Flaveria* leaves. The numbers of differentially expressed genes in each section of the leaf are depicted to the left (up in both C_3_ *F. pringlei* and *F. robusta*) and to the right (up in both C_4_ *F. bidentis* and *F. trinervia*) of the leaf schematic. **B.** Principal Components Analysis of the differentially expressed genes in all four species shows that the first dimension is associated with leaf maturation, but the second dimension is associated with photosynthetic pathway. C_3_ species are depicted in red, C_4_ species in blue, and each stage is numbered from 1 (base) to 6 (tip). **C.** Summary of MapMan categories associated with the top decile of differentially expressed genes in each species. *Fp* = *Flaveria pringlei*, *Fr* = *Flaveria robusta*, *Fb* = *Flaveria bidentis* and *Ft* = *Flaveria trinervia*.

### Kranz anatomy during leaf maturation

Using cleared leaves, veins were traced to visualise their development. In the most basal part of the leaf, major veins were present but in the two C_4_ species, the highest order veins were still being laid down. Some of these developing higher order veins could be detected by the specific patterns of periclinal cell divisions associated with their production (Supplementary Figure 6). Vein density was determined in each of the six sections along the leaf developmental gradient, and this showed markedly different developmental patterns in the C_3_ and C_4_ pairs of *Flaveria*. (Figure 5A). Vein density was lower at the base in C_4_ compared with C_3_ leaves of *Flaveria*, but during the transition from the base to the middle of the leaf density increased dramatically in the C_4_ species (Figure 5B). In both C_3_ and C_4_ pairs, vein density then decreased from the middle to the tip of the leaf. The reduction in vein density as leaves matured is likely associated with expansion of M and BS cells. In the two most basal sections of the leaf cell division was still ongoing and procambial strands were initiated first with a periclinal and then an anticlinal division, to derive BS cells (Supplementary Figure 6). These basal portions of the leaf therefore appear most interesting to interrogate for candidate genes associated with these processes. The *de novo* assembled transcriptomes from the four *Flaveria* species were therefore analysed for homologues of genes known to impact on vein formation in others species.

**Figure 5:**
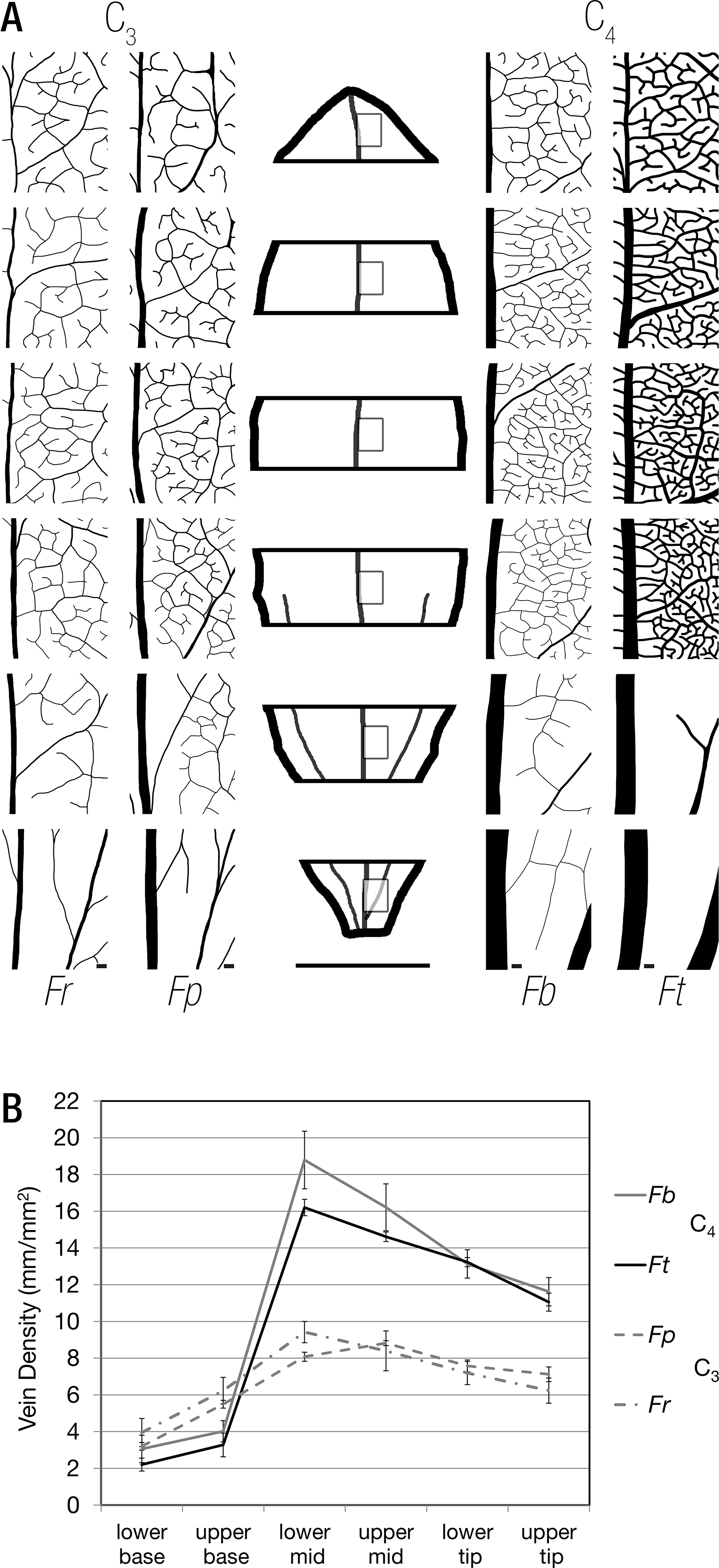
Vein density increases dramatically between the upper base and lower mid leaf sections of the C_4_ species compared with the C_3_ species. Representative vein traces from base to tip of all four species illustrating the maturation of vein density. *Fp = Flaveria pringlei, Fr = F. robusta, Fb = Flaveria bidentis and Ft = Flaveria trinervia*. The leaf outline in the centre indicates that vein density measurements were obtained from the centre of each section to the right of the midrib. **B.** Quantitation of vein density in each section for the four species. In all species, vein density was highest in the middle and decreased towards the tip of leaves. A rapid increase in vein density occurs between the upper base and lower mid in both C_4_ species. Error bars represent one standard error of the mean. Scale bars represent 100μm for the vein traces and 0.5cm for the leaf outline.

Six genes that effect vein formation in *A. thaliana* showed different behaviours in the C_4_ *Flaverias* compared with the C_3_ species (Figure 6). In all cases absolute transcript abundance tended to be higher at the base of the C_4_ leaves compared with the base of the C_3_ species. However, the most consistent differences in expression were found for *Arabidopsis thaliana HOMEOBOX-GENE-8* (*ATHB8*). *SHR* and *SCR* have been implicated with the development of C_4_ Kranz anatomy (Slewinski *et al.*, 2012; Slewinski, 2013). *SHR* was detected in three of the species with a descending pattern but no clear difference between C_3_ and C_4_. *SCR* also showed a descending pattern but again with no clear difference between photosynthetic types. Of Scarecrow-like genes, only *SCARECROW-LIKE 14* was clearly higher in C_4_ and showed a parabolic expression pattern, being most highly expressed at the base and the tip (Supplementary Data 3). Transcripts encoding the auxin response factors ARF3, ARF8 and also IAA7 showed differences in the timing of expression in C_4_ compared with C_3_ species (Supplementary Data 3).

**Figure 6:**
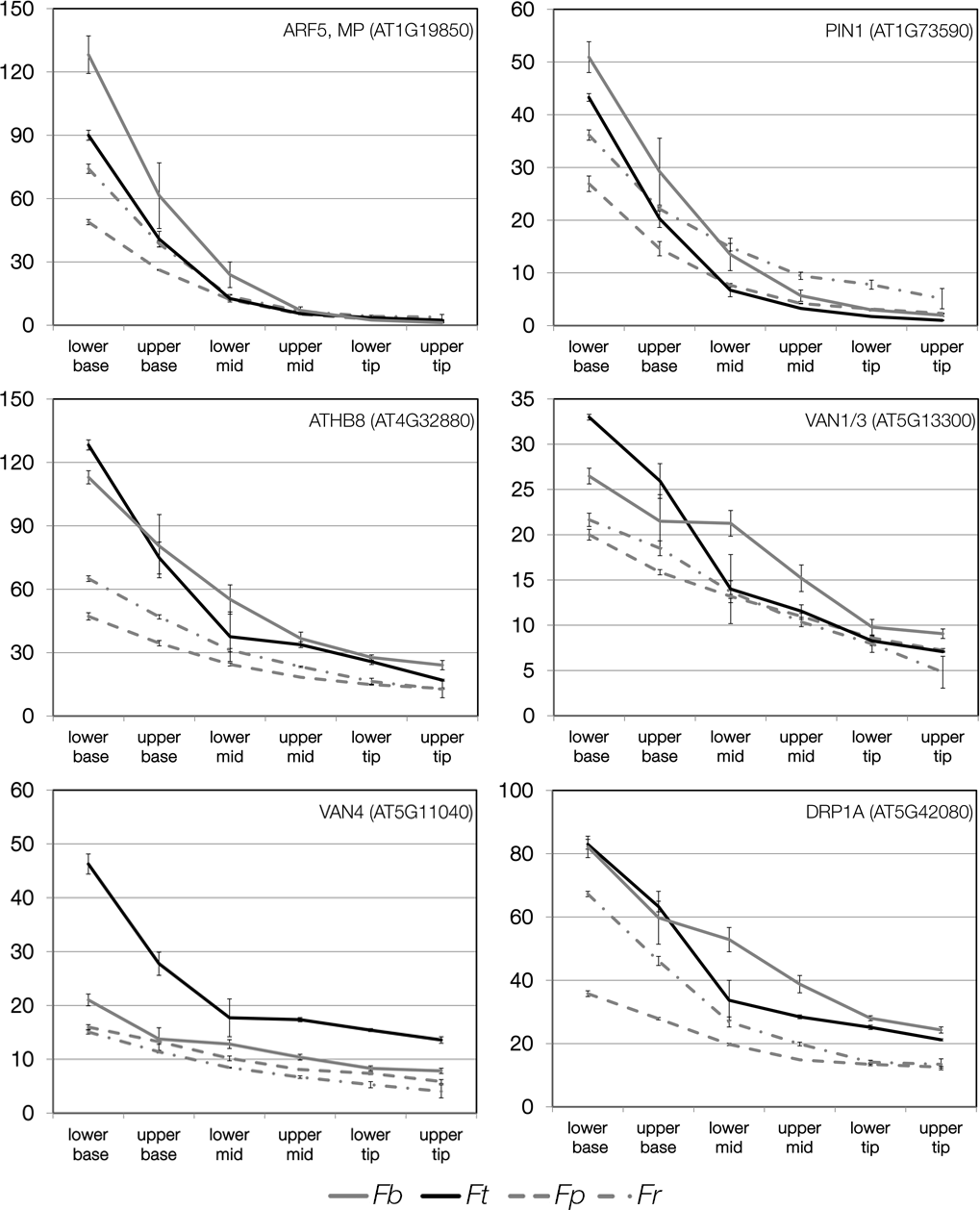
Patterns of transcript abundance in the four *Flaveria* species for genes known to be involved in vein formation. Transcript abundance is shown in transcripts per million (TPM), and the parts of the leaf are annotated on the x-axis. The two C_4_ species are shown with solid lines, and the two C_3_ species with dashed lines. *Fp = Flaveria pringlei, Fr = Flaveria robusta, Fb = Flaveria bidentis and Ft = Flaveria trinervia*.

The same six regions from leaf base to tip used to define vein density were next used to quantify maturation of BS and M cells (Figure 7). Leaf thickness was higher in the C_3_ compared with the C_4_ species, and leaf expansion continued for longer in the C_3_ leaves (Figure 7A). The BS in the C_3_ species was less regular than in the C_4_ species, and individual cell size within the BS was more variable (Figure 7A). However, it was notable that the cross-sectional area of BS cells in the C_3_ species was larger than that of C_4_ species, particularly towards the tip. It has been proposed that a large BS cell size is a key early event associated with the evolution of C_4_ photosynthesis (Williams *et al.*, 2013; Christin *et al.*, 2013; Griffiths *et al.*, 2013), and so it appears that within *Flaveria* this is a pre-existing trait. It was noticeable that the cross-sectional area of M cells from the two C_3_ species was around five times greater than M cells of the C_4_ species (Figure 7B). This strongly implies that compared with C_3_ species, the reduced leaf depth of C_4_ species is associated with an inhibition of M cell expansion as well as a reduced number of cell layers. A number of genes that have been annotated as having roles in cell proliferation or expansion showed behaviours that may explain the reduced cell expansion in leaves of both C_4_ species. Most notable was *DWF4*, a 22α-hydroxylase that catalyses the rate-limiting step of Brassinosteroid synthesis and so controls cell expansion. In both C_3_ species, *DWF4* transcript abundance increased along the leaf gradient but in the C_4_ species its transcripts were barely detectable (Supplementary Data 4). These data imply that *DWF4* is a strong candidate for the reduced expansion associated with maturation of the C_4_ leaf.

**Figure 7:**
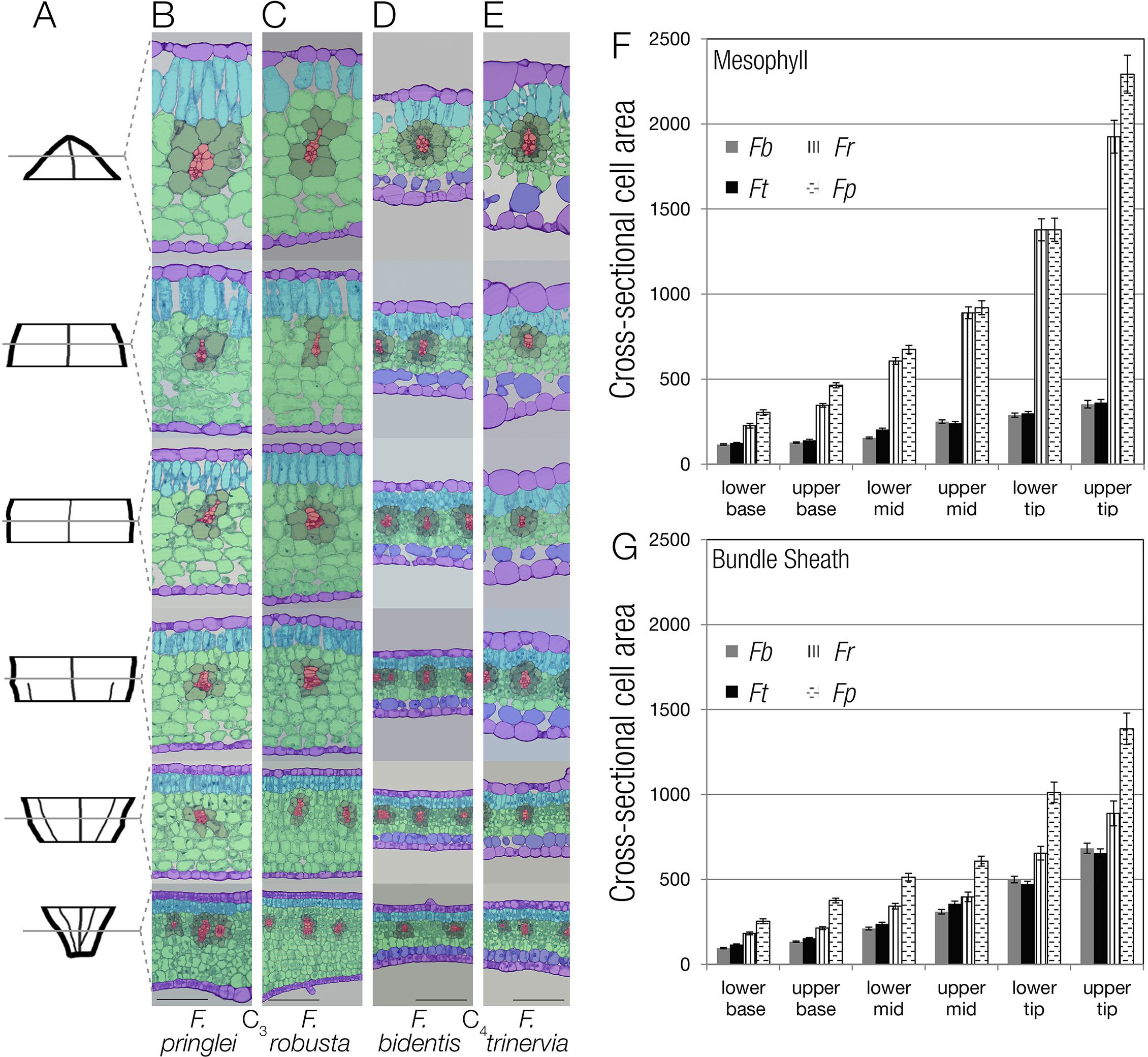
During leaf maturation mesophyll cell expansion is reduced in the C_4_ compared with the C_3_ species. **A**. Representative leaf outline illustrating leaf sampling. **B-E**. Representative transverse sections from base to tip of leaves from C_3_ *F. pringlei* (**B**) and *F. robusta* (**C**), as well as *F. bidentis* (**D**) and *F. trinervia* (**E**). **F&G.** Quantification of cross-sectional cell areas for mesophyll (**F**) and bundle sheath (**G**) cells along leaf maturation gradient. Each cell is false coloured-coded: veins (red), bundle sheath (dark green), spongy mesophyll (light green), palisade mesophyll (turquoise), parenchyma layer with no or few chloroplasts (blue) and epidermis (lilac). Species abbreviations are as follows: *Flaveria pringlei (Fp), Flaveria robusta (Fr), Flaveria bidentis (Fb), Flaveria trinervia, (Ft)*. Scale bars (**B-E**) represent 100μm.

A prolonged rate of cell division in the C_4_ *Flaveria* species is also supported by transcript profiles. A number of genes implicated in cell division are higher in basal sections of C_4_ than C_3_ leaves including *PLE* (Müller *et al.*, 2004), *MOR1* (Whittington *et al.*, 2001), *ATK1* (Marcus *et al.*, 2003) and *ATK3* (Mitsui *et al.*, 1994) which are involved in microtubule organisation, and *SCC3* (Chelysheva *et al.*, 2005) and *SYN1* (Cai *et al.*, 2003) which are implicated in sister chromatid pairing (Supplementary Data 4). Transcripts derived from genes associated with DNA replication and repair including *SOG1* (Yoshiyama *et al.*, 2009), *TOP6B* (Gilkerson and Callis, 2014), *MutS* homologs *MSH* (*MSH3, MSH4, MSH7*) (Culligan and Hays, 2000) and multiple components of the Mini-chromosome maintenance (MCM) complex (*MCM2, MCM3, MCM5, MCM6*) (Tuteja *et al.*, 2011) were also more abundant in the C_4_ species at the base of the leaf (Supplementary Data 4). Good candidate regulators of cell division include MYB3R4 which forms a complex with E2FB and *RETINOBLASTOMA-RELATED PROTEIN* (RBR1) to activate cell-cycle progression (Haga *et al.*, 2011), and *DEL1* which maintains cell division by repressing entry into the endocycle (Vlieghe *et al.*, 2005), both of which are higher in the base of C_4_ leaves and maintain expression longer into the leaf gradient.

A reduction along the leaf gradient in the abundance of transcripts encoding CPP SYNTHASE 1 (CPS1), which catalyses the first step of gibberellin synthesis, and a spike in transcripts of *GA2ox2* (Supplementary Data 4), which degrades gibberellin, between upper base and lower mid in C_4_ leaves, is in keeping with findings that a decrease in GA levels controls the transition between cell division and expansion in Maize (Nelissen *et al.*, 2012). A corresponding pattern cannot be seen in C_3_ leaves, which may mean that the transition between cell division had already occurred at the base of C_3_ lineages, or is mediated by another mechanism.

### Chloroplast maturation within the leaf gradient

Species that primarily use NADP-ME to decarboxylate malate in the C_4_ BS commonly develop dimorphic chloroplasts with M cells containing stacked thylakoids (grana) and the BS cells containing fewer grana (Edwards and Walker, 1983; Edwards and Voznesenskaya, 2011). To provide insight into the dynamics of chloroplast development and maturation in C_3_ and C_4_ *Flaveria* species, transmission electron microscopy was used to investigate chloroplast structure in each of the six stages along the leaf gradient. Dimorphic chloroplasts were observed in the C_4_ species but not in the C_3_ species (Figure 8A-D). Towards the base of the C_4_ leaves, both BS and M chloroplasts contained 2-3 granal lamellae per stack (Supplementary Figure 7). In more mature parts of the leaf levels of stacking were reduced in BS cells but increased in M cells (Figure 8A-D). In both C_3_ species, a developmental gradient in granal stacking was also visible from base to tip, with more mature parts of the leaf showing increased levels of granal stacking in both cell types (Supplementary Figure 7). Thus, a key difference between the C_3_ and C_4_ leaves was the reduction in granal stacking in C_4_ BS cells from the middle of the leaf.

**Figure 8:**
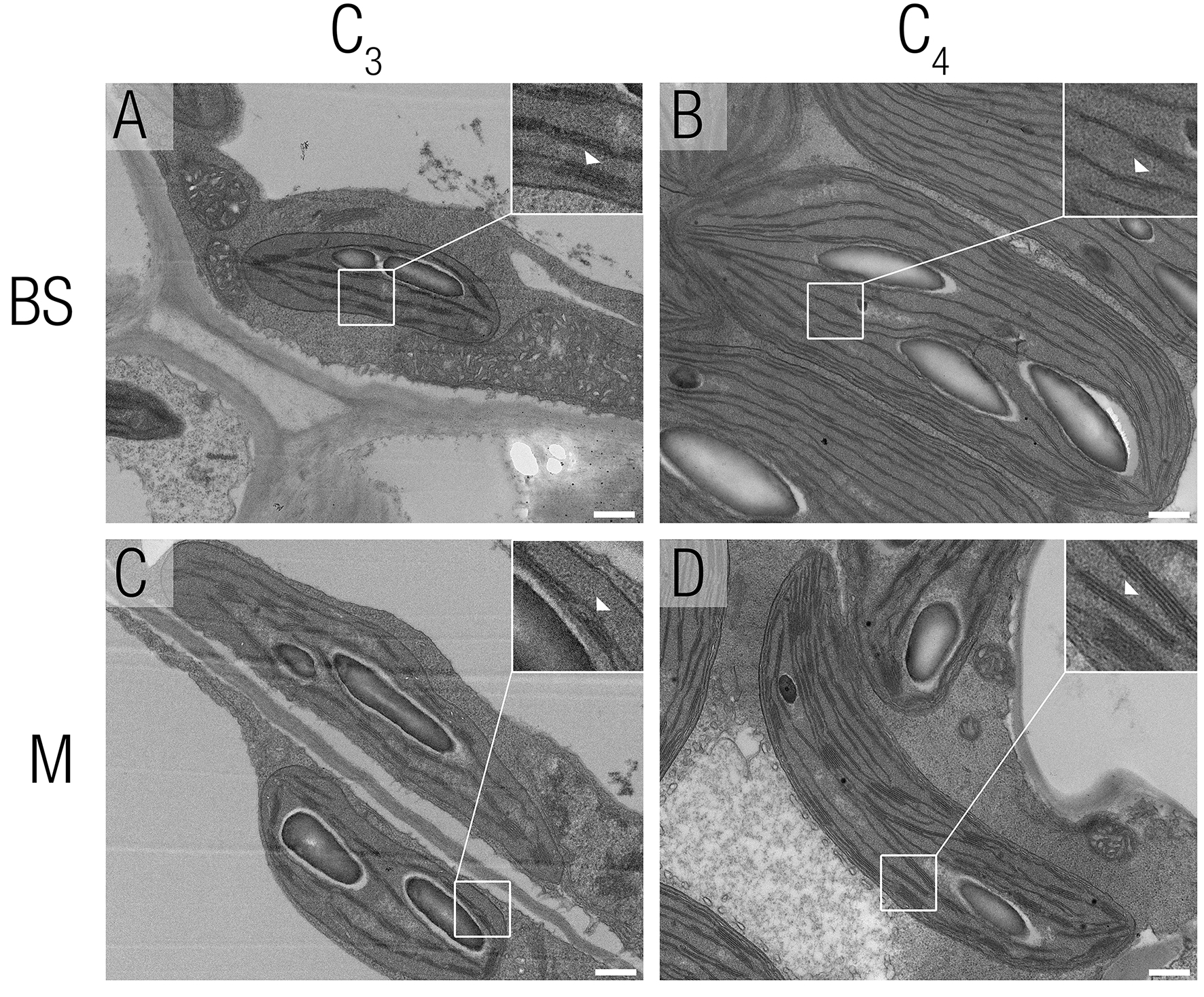
Chloroplast ultrastructure from mesophyll and bundle sheath cells from C_3_ *F. pringlei* and C_4_ *F. bidentis*. Transmission electron microscopy was used to investigate chloroplast ultrastructure. **A&B,** Bundle sheath chloroplasts from C_3_ *Flaveria pringlei* and C_4_ *Flaveria bidentis*. **C&D,** Mesophyll chloroplasts from C_3_ *F. pringlei* and C_4_ *F. bidentis*. Little granal stacking was seen in bundle sheath chloroplasts from *F. bidentis* (**B**) whereas it could be observed in C_3_ bundle sheath and in all mesophyll chloroplasts. Insets show close-up of thylakoids with white arrowheads indicating thylakoid stacks (**A**, **C**, **D**) or the lack of extensive stacking (**B**). Scale bars represent 500nm.

The control of photosynthesis gene expression and chloroplast development has previously been linked to a pair of transcription factors known as GOLDEN-LIKE 1 and 2 (GLK1 and GLK2) (Supplementary Data 5). Unfortunately, *de novo* assembly did not generate a complete set of contigs for either GLK1 or GLK2 from all four species so we were unable to determine if their abundance correlated with the observed changes in chloroplast development. However, other candidate genes that can be associated with chloroplast maturation were identified. For example, cytokinin is known to play a role in chloroplast maturation, and increased abundance of transcripts encoding the cytokinin receptors AHK2 and AHK3, along with the downstream effector ARR1 in C_4_ species suggests that enhanced cytokinin signalling may play a role in these alterations to the C_4_ chloroplast. Additionally, it was also notable that the behaviour of two nuclear encoded chloroplast RNA polymerases (RpoTp and RpoTmp) (Swiatecka-Hagenbruch *et al.*, 2008) showed clear differences between the C_3_ and C_4_ *Flaveria* species. Transcripts encoding RpoTp, which controls *ycf1* that is involved in plastid protein import were very low in both C_4_ species. Further indications to changes in chloroplast protein import related to components of the TIC/TOC (Supplementary Data 5). Transcripts encoding the outer envelope receptor proteins TOC34 and TOC159 were higher in the base and peaked later in the C_4_ than the C_3_ species. Additionally, TOC75, the major channel constituent for protein import into the chloroplast (Li and Chiu, 2010) was lower at the base of C_4_ leaves. These data would be consistent with protein import playing a role in the establishment of dimorphic chloroplasts. Chloroplast maturation also correlated with increased abundance of nuclear-encoded chloroplast sigma factors, which direct the nuclear-encoded chloroplast RNA polymerase (NEP) to promoters.

Previous studies have demonstrated that C_4_ plants alter Photosystem activity to adjust ADP/ATP and NADPH ratios to ensure proper functioning of the carbon pump. Consistent with this was the C_4_ specific up-regulation of SIG3, which initiates transcription of *psbN* (Zghidi *et al.*, 2007) which is involved in repair of PSII complexes (Torabi *et al.*, 2014). It was also notable that transcripts encoding components of the NDH complex (*NDF1, NDF6* and *PIFI*), as well as genes involved in cyclic electron transport (*PGR5* and *PGRL1A*) were more abundant in both C_4_ species, and these differences became more apparent from base to tip (Supplementary Data 5). Chloroplast dimorphism also coincided with an increase in transcript abundance of *CRR1*, which is proposed to be involved in NDH complex assembly.

Analysis of electron micrographs also revealed that chloroplast length in the BS increased from base to tip in the C_4_ species more than in the C_3_ species (Figure 9). Two members of the *REDUCED CHLOROPLAST COVERAGE* (*REC*) gene family, REC2 and REC3, knockout of which causes a reduction in chloroplast size (Larkin *et al.*, 2016) were up-regulated in the C_4_ species. Further, *FtsZ1-1* which is involved in chloroplast division, was significantly down-regulated in the C_4_ compared with the C_3_ species (Supplementary Data 5). As reduced division of chloroplasts leads to increased size (Pyke and Leech, 1994; Pyke, 1997) the reduced expression of chloroplast division genes in C_4_ plants could lead to their increased size in BS cells.

**Figure 9:**
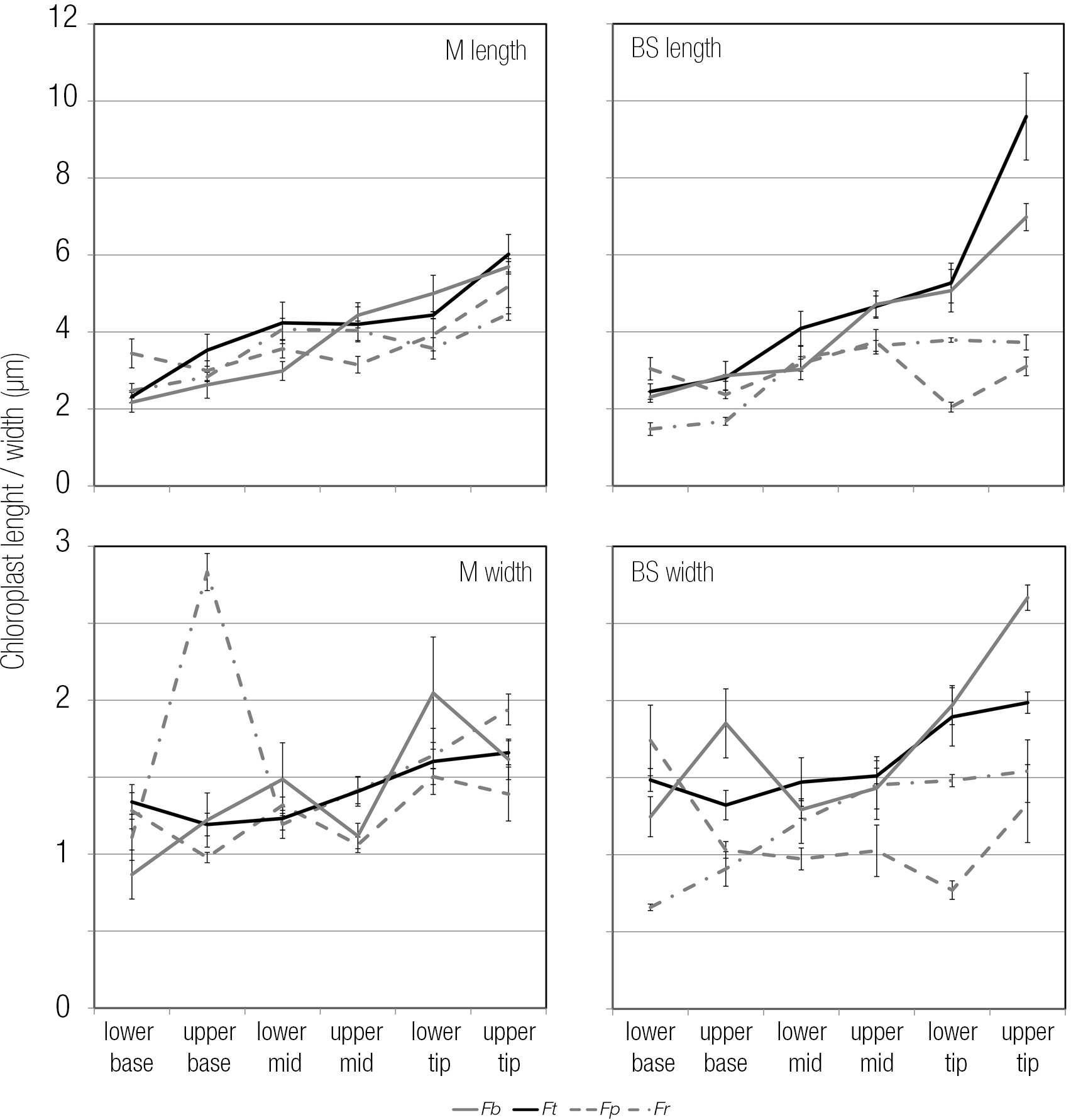
Quantitation of chloroplast length and width from mesophyll and bundle sheath cells. Length and width were taken from thin sections used in transmission electron microscopy. For each species five chloroplasts were measured per stage and cell type. Error bars represent one standard error of the mean. The two C_4_ species are shown with solid lines, and the two C_3_ species with dashed lines.

### Convergence in transcript abundance between independent C_4_ lineages

By combining our data from the four *Flaveria* species with publically available datasets, we next sought to investigate whether separate lineages of C_4_ plants shared common changes in transcript abundance compared with their C_3_ congeners. A recent study compared transcriptomes derived from leaf developmental gradients of *Tarenaya hassleriana* (C_3_) and *Gynandropsis gynandra* (C_4_) from the Cleomaceae (Külahoglu *et al.*, 2014). In contrast to our work, whole leaves were staged by age (Külahoglu *et al.*, 2014) and changes in transcript abundance compared with non-photosynthetic tissues such as roots. This analysis identified a set of thirty-three genes that could be annotated with unique *A. thaliana* identifiers. This set of genes showed root expression in the C_3_ species *T. hassleriana* but leaf expression in the C_4_ species *G. gynandra* (Külahoglu *et al.*, 2014), and so it was proposed that these were key genes that neofunctionalise to become involved in C_4_ photosynthesis. Of these thirty-three genes, homologues to fifteen were detected in all four *Flaveria* species, but only three showed up-regulation in the C_4_ compared with C_3_ species in our study and three showed higher expression in the two C_3_ species (Supplementary Data 6).

Previously, transcript abundance in developing leaves of the C_4_ dicot *Gynandropsis gynandra* (Aubry *et al.*, 2014) has been compared with the C_4_ monocot maize (Li *et al.*, 2010; Pick *et al.*, 2011). Despite the very wide phylogenetic distance between these species, eighteen transcription factors that showed the same patterns of behaviour in both C_4_ *G. gynandra* and C_4_ maize were identified (Aubry *et al.*, 2014). Ten of these eighteen candidates were also detected in all four *Flaveria* species. Five of those showed ascending behaviour and higher transcript abundance in the two C_4_ species. These were RAD-LIKE6 (RL6, AT1G75250); AT2G05160, a CCCH-type zing finger family protein with RNA-binding domain; AT2G21530, a SMAD/FHA domain containing protein; RNA Polymerase Sigma Subunit C (SIGC, AT3G53920) and BEL1-Like Homeodomain1 (BLH1, AT5G67030) (Supplementary Data 7). The five genes have therefore been shown to exhibit the same behaviour along leaf developmental gradients in C_4_ species in the monocots and dicots (both rosids (Cleomaceae) and asterids (Asteraceae)). This strongly supports the notion that independent lineages of C_4_ plant have recruited homologous *trans*-factors to induce the C_4_ system.

## Summary and Conclusions

Most previous analysis derived from comparisons of RNA-seq of C_3_ and C_4_ species have either relied on sampling one con-generic C_3_ and C_4_ species pair, or analysis of mature leaf tissues. The number of genes that are differentially expressed between any two species can be very high simply due to species-specific differences, which do not relate to their photosynthetic pathway. In this work we identify genes that are consistently differentially expressed in multiple C_3_ and C_4_ species from the same genus along a leaf maturation gradient. In the *Flaveria* genus, the dataset therefore indicates that ~200 genes are consistently up-regulated in mature tissue using C_4_ compared with C_3_ photosynthesis. In young tissue that is still undergoing differentiation, the number of genes up-regulated in the C_4_ species is around double this number. Leaf maturation and then photosynthetic pathway were responsible for the greatest amount of variation in transcript abundance. Overall, the analysis therefore provides quantitation of changes in gene expression associated with C_3_ and C_4_ photosynthesis, candidate genes underlying the alterations in leaf characteristics associated with these pathways, and through comparison with independent C_4_ lineages outside of the Asteraceae, insight into the extent to which parallel evolution underlies the convergent and complex C_4_ trait.

## Supplementary Data

**Supplementary Figure 1:** Mature leaf vein density and cell size.

**Supplementary Figure 2:** Leaf anatomy maturation gradient in C_3_ and C_4_ *Flaveria.*

**Supplementary Figure 3:** RNA quality from sections of each species.

**Supplementary Figure 4:** TPM correlation matrix.

**Supplementary Figure 5:** GO term analysis presented as heatmaps for each *Flaveria* species.

**Supplementary Figure 6:** Cell division and vein development in the base samples of *Flaveria*.

**Supplementary Figure 7:** Chloroplast maturation in mesophyll and bundle sheath cells for each species.

**Supplementary Data 1:** TPM values for all four species for all stages and all replicates.

**Supplementary Data 2:** Summary of transcripts that were upregulated in both C_3_ species or both C_4_ species at each stage.

**Supplementary Data 3:** Transcript abundance of *SHR, SCR* and *SCR-like* genes as well as auxin response genes.

**Supplementary Data 4:** Transcript abundance of genes associated with cell division.

**Supplementary Data 5:** Transcript abundance of genes associated with chloroplast maturation and development.

**Supplementary Data 6:** Comparison of data derived from this study with genes identified by Külahoglu et al (2014).

**Supplementary Data 7:** Comparison of data derived from this study with genes identified by Aubry et al (2014).

**Supplementary Table 1:** Number of reads and transcripts, per stage and species.

## Acknowledgements

Seeds for all *Flaveria* species used were kindly provided by Udo Gowik (Universität Düsseldorf). European Union *3to4* project and the Biotechnology and Biological Sciences Research Council (BBSRC) grant BB/J011754/1 funded the project.

## Author contributions

BMCK and JMH designed the study. BMCK grew the plants, undertook the anatomical analysis and isolated the RNA; RS-U and CB undertook the *de novo* transcriptome assembly and annotation of contigs; BMCJ, SJB and IR-L analysed the RNA-seq data; BMCK, SJB and JMH wrote the paper.

